# ULK-mediated phosphorylation of ATG16L1 promotes xenophagy, but destabilizes the ATG16L1 Crohn’s mutant

**DOI:** 10.1101/387563

**Authors:** Reham M. Alsaadi, Truc T. Losier, Wensheng Tian, Zhihao Guo, Ryan C. Russell

**Author notes:** denotes equal contribution (Reham M. Alsaadi, Truc T. Losier). Address for correspondence: 451 Smyth Rd, Ottawa, Ontario, K1H 8M5, Canada.

## Abstract

Autophagy is a highly regulated catabolic pathway that is potently induced by stressors including starvation and infection. An essential component of the autophagy pathway is an ATG16L1-containing E3-like enzyme, which is responsible for lipidating LC3B and driving autophagosome formation.

ATG16L1 polymorphisms have been linked to the development of Crohn’s disease (CD) and phosphorylation of CD-associated ATG16L1 (caATG16L1) has been hypothesized to contribute to cleavage and autophagy dysfunction. Here we show that ULK1 kinase directly phosphorylates ATG16L1 in response to infection and starvation. Moreover, we show that ULK-mediated phosphorylation drives the destabilization of caATG16L1 in response to stress. Additionally, we found that phosphorylated ATG16L1 was specifically localized to the site of internalized bacteria indicating a role for ATG16L1 in the promotion of anti-bacterial autophagy. Lastly, we show that stable cell lines harbouring a phospho-dead mutant of ATG16L1 have impaired xenophagy. In summary, our results show that ATG16L1 is a novel target of ULK1 kinase and that ULK-signalling to ATG16L1 is a double-edged sword, enhancing function of the wildtype ATG16L1, but promoting degradation of caATG16L1.

## Introduction

Macroautophagy (hereafter referred to as autophagy) is a cellular degradative process capable of degrading a vast array of substrates including cytoplasm, organelles, aggregated macromolecules, and pathogen^1^. Autophagic cargo is first sequestered by the *de novo* formation a double-membraned vesicle called an autophagosome, which matures into a degradative vesicle after fusion with lysosomes.

Autophagosome formation is driven by a set of autophagy-related (ATG) genes, which include a protein kinase (Unc 51-like kinase 1; ULK1), a lipid kinase (vacuolar protein sorting 34; VPS34), and a trimeric E3-like enzyme (ATG5-ATG12/ATG16L1)^1^. These enzymes are all required for autophagy initiation and are tightly regulated by upstream stress-sensitive signalling. One of the best characterized upstream regulators of the autophagy pathway is mTORC1, which potently inhibits autophagy induction through direct phosphorylation of the ULK1 and VPS34 kinase complexes^2–5^. mTORC1 activity is repressed, thereby allowing autophagy induction, in response to a myriad of stressors including nutrient or cytokine starvation, reactive oxygen species, or infection^6–8^.

Mammals have two homologues of the yeast ATG1, ULK1 and ULK2, which are functionally redundant for autophagy induction (ULK1 and ULK2 will collectively be referred to as ULK hereafter)^9^. Under basal conditions, mTORC1-mediated phosphorylation represses ULK activity; however, starvation releases this inhibitory phosphorylation and upregulates ULK^2^. Activated ULK then phosphorylates several components of the pro-autophagic ATG14-containing VPS34 complexes^10–12^. Autophagic VPS34 complexes are recruited to the phagophore where they phosphorylate phosphatidylinositol (PtdIns) to produce phosphatidylinositol(3)phosphate (PtdIns(3)P)^13^. PtdIns(3)P functions as a platform bridging downstream components to promote autophagosome formation. Additionally, mTORC1 has been shown to directly mediate the activity of VPS34 complexes, thereby allowing a tight regulation of autophagy initiation in response to stresses^3^. Downstream of VPS34, ATG16L1 forms a trimeric complex with ATG5 and ATG12. ATG16L1 is the subunit responsible for recruiting the E3-like enzyme to the phagophore^1,14^. ATG12 acts to recruit microtubule-associated protein 1 light chain 3 (LC3) to the expanding autophagosomal membrane and ATG5 catalyzes the lipidation of this substrate, thereby driving autophagosome formation.

Activation of anti-bacterial autophagy (hereafter referred to as xenophagy) involves these 3-key enzymes in the autophagy pathway, but also requires xenophagy-specific proteins involved in pathogen-sensing that signal to the autophagy machinery during infection^8^. For instance, galectin-8 detects damaged *Salmonella-*containing vacuoles (SCV) and subsequently activates xenophagy through recruitment of the autophagy receptor NDP52^15^. Immunity related GTPase M (IRGM) has been shown to act as a scaffold bringing together ULK, Beclin-1-containing VPS34 complexes, and ATG16L1 to promote xenophagy initiation^16^. In addition to IRGM, ATG16L1-containing enzyme is also regulated by activation of intracellular (NOD2) sensors of bacterial peptidoglycan, where NOD2 binds ATG16L1 recruiting the LC3-lipidating enzyme to the site of bacterial infection^17^.

Interestingly, several of the proteins involved in xenophagy induction (ATG16L1 and IRGM) and pathogen detection (NOD2 and TLR4) have been linked to Crohn’s disease (CD), but are not found in the related chronic inflammatory bowel disease ulcerative colitis (UC)^18^. Genome-wide association studies have linked a non-synonymous single nucleotide polymorphism (SNP) in ATG16L1 that substitutes threonine 300 for alanine with an increased susceptibility for CD^19^. Molecular characterization of the CD-associated ATG16L1 (caATG16L1) has shown that stresses such as starvation or pathogen infection enhance the susceptibility of caATG16L1 to caspase-mediated cleavage^20–23^. Enhanced cleavage of caATG16L1 has been shown to lead to an increase in inflammatory cytokine secretion and a decrease in xenophagy, which are thought to contribute to CD^21,24-26^. Interestingly, a recent study has found that IκB kinase subunit IKKα is capable of phosphorylating ATG16L1 on Serine 278 (S278), which regulates the sensitivity of caATG16L1 to caspase cleavage^24^. The caspase cleavage site on ATG16L1 lies in between the S278 phosphorylation site and the T300A Crohn’s SNP. This raises the interesting possibility that phosphorylation of ATG16L1 in response to infection leads to inappropriate cleavage if the site is in close proximity to the T300A mutation. ATG16L1 contains several conserved serine/threonine residues proximal to T300, which may also be phosphorylated and may potentially regulate ATG16L1 function.

However, it remains to be seen what effect phosphorylation has on wild-type ATG16L1 and if other stressors or kinases regulate ATG16L1 phosphorylation.

## Results

Starvation has been described to trigger caspase-mediated cleavage of ATG16L1 containing a common amino acid substitution (T300A)^21^. However, IKKα has not been implicated in starvation-induced autophagy. Therefore, we hypothesized that ULK, the only protein kinase in the autophagy pathway, may phosphorylate ATG16L1 under starvation. To test this hypothesis we performed an *in vitro* kinase assay using either purified ULK1 or ULK2 with recombinant ATG16L1 as substrate.

Interestingly, we found that both ULK1 and ULK2 were capable of phosphorylating ATG16L1 *in vitro* (Fig. 1A). In order to narrow down the site of phosphorylation we repeated the kinase assay using truncations of ATG16L1. We found that the truncation mutant lacking amino acids 254-294 was a very poor substrate for ULK, indicating that the primary site(s) of ULK-mediated phosphorylation are located in this region (Fig. 1B). Amino acids 254-294 are serine/threonine rich, containing 10 conserved residues (Fig. 1C). Therefore, to identify the residue(s) that are phosphorylated by ULK in this region we repeated the kinase assay on full length ATG16L1 and performed mass spectrometry analysis. Our results revealed a single high confidence phosphorylation site on serine 278 (Fig. S1 and marked in green in Fig. 1C) and another of slightly lower confidence on serine 287 (Fig. S1 and marked in grey in Fig. 1C), both of which map to the region of ATG16L1 we previously identified as required for ULK-mediated phosphorylation (Fig. 1B). Peptide coverage in the mass spectrometry was 80% across the whole protein and only two S/T residues were missed in the putative 254-294 region. To confirm the major site(s) of phosphorylation on ATG16L1 we mutated S278 and S287 singly in the full length protein and performed another *in vitro* ULK kinase assay. Interestingly, we observed a nearly complete loss of ULK-mediated phosphorylation in the S278 mutant, which was comparable to the reduction in the 254-294 truncation mutant (Fig. 1D). This indicates that the major site of phosphorylation on ATG16L1 is S278, which is the same residue previously identified as a site for IKKα-mediated phosphorylation^24^. Next, we created a phospho-specific antibody against S278 of ATG16L1 and tested its specificity by co-transfection of wild-type or mutant ULK and ATG16L1. Excitingly, we observed that ULK phosphorylates ATG16L1 in cells and that our antibody was specific to the phosphorylated form of the protein with little to no signal against ATG16L1 (278A) or wild-type ATG16L1 cotransfected with kinase-dead ULK (Fig. 1E). Collectively, these results show that ATG16L1 is a direct target of ULK and that the primary site of phosphorylation is S278.

**Figure 1.**
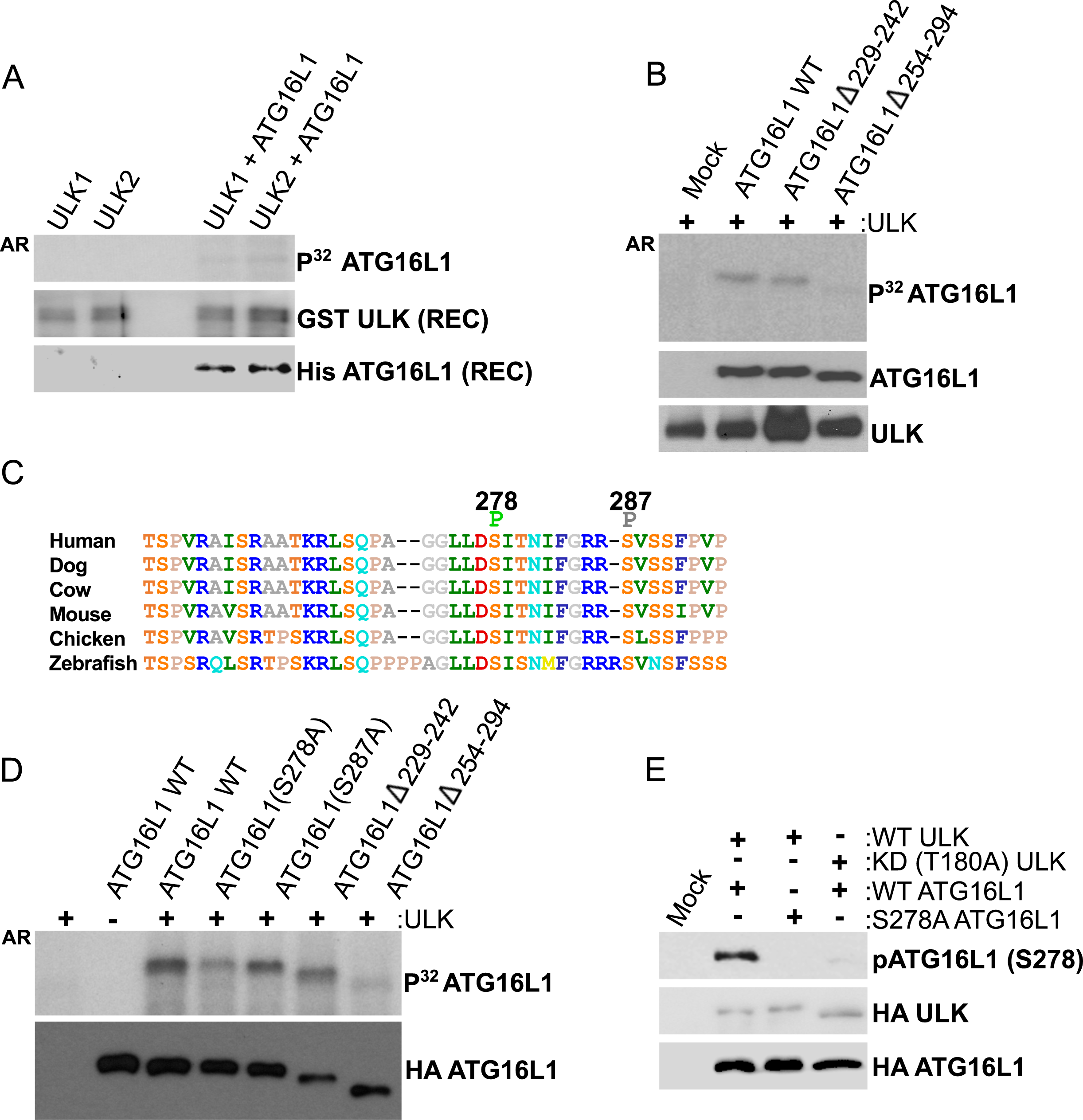
ATG16L1 is phosphorylated by ULK. **(A)** *in vitro* kinase assays were performed using purified recombinant kinases (ULK1 and ULK2) and substrate (ATG16L1) in the presence of radiolabelled ATP. ULK and ATG16L1 inputs were examined by Western blot (WB) and substrate phosphorylation was analyzed by autoradiography (AR). **(B)** Full-length or truncated versions of ATG16L1 were subjected to an *in vitro* ULK kinase assay. ULK1 and ATG16L1 inputs were examined by Western blot and target phosphorylation by autoradiography. **(C)** ATG16L1 was phosphorylated in an *in vitro* ULK kinase reaction and analysed by mass spectrometry. Phosphorylation of S278 and S287 in human (S278 marked in green, S287 marked in grey) was identified with high and low confidence, respectively. Conservation of amino acids 254-294 are shown using the Shapely colour scheme. Mass spectrometry was performed on a single experiment. **(D)** Full-length, truncated, or mutated HA-ATG16L 1 was purified from mammalian cells and subjected to an *in vitro* ULK kinase assay. ATG16L1 inputs were analysed by WB and target phosphorylation by AR. **(E)** HEK293A cells were transfected with wild-type or phospho-dead ATG16L1 in the presence of wild-type or kinase-dead ULK. The specificity of phospho antibody and inputs were examined by WB.

We next sought to determine if ULK regulated ATG16L1 phosphorylation endogenously and whether this signalling was responsive to starvation. ULK wild-type or ULK1/2 double knockout (dKO) cells were starved for amino acids, either with amino acid-free DMEM or HBSS, followed by analysis of pATG16L1 levels by Western blot of whole cell extracts. Starvation potently inhibits mTORC1-signalling, as demonstrated by loss of S6K phosphorylation, which is a prerequisite for ULK activation (Fig. 2A).

**Figure 2.**
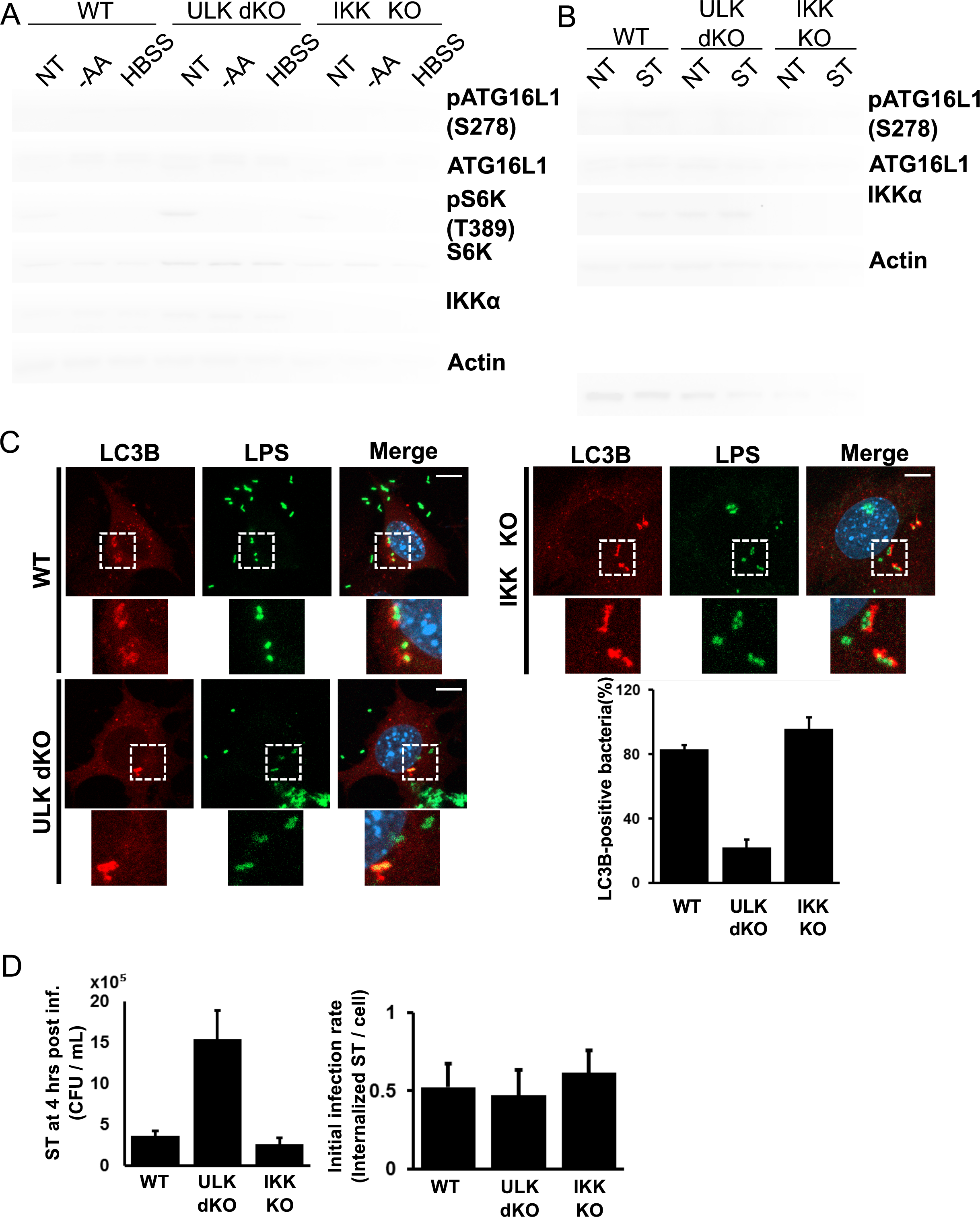
ULK is required for phosphorylation of ATG16L 1 and xenophagy induction. **(A)** Wild-type, ULK double knockout (dKO), or IKK KO mouse embryonic fibroblasts (MEFs) were incubated with either complete medium, amino acid-deficient DMEM, or HBSS for 1 hour. Samples were immunoblotted using the indicated antibodies. **(B)** Wild-type, ULK dKO, or IKK KO MEFs were infected with log phase Salmonella for 2 hours; bacteria-containing media was then removed and cells were incubated with gentamycin (50µg/ml)-containing DMEM for 2 hours. Samples were immunoblotted using the indicated antibodies. **(C)** Wild-type, ULK dKO, or IKK KO MEFs cells were infected with Salmonella for 1 hour in the presence of Bafilomycin A 1. Bacteria were stained using anti-LPS antibodies to analyze localization in addition to the autophagy marker LC3B. Representative immunofluorescent images of LC3B and LPS are shown in the left panel (scale bars, 1 0µm). Quantification of bacteria undergoing autophagic clearance from 8 fields of view from a representative experiment is shown in the right panel. **(D)** Wild-type, ULK dKO and IKKa KO MEFs were infected with Salmonella for 1 hour. Xenophagy rates were examined through Colony Forming Unit (CFU) assays. Quantification of infection rates by immunofluorescence is demonstrated in the right panel.

Importantly, we observed that starvation resulted in a clear increase in endogenous ATG16L1 phosphorylation only in cells containing ULK (Fig. 2A, lanes 1-6). Notably, our phospho-antibody only recognizes the slower migrating ATG16L1α isoform and is observed as a single band. As IKKα was previously described to phosphorylate ATG16L1 on S278 under infection we also tested the requirement for IKKα in starvation-induced ATG16L1 phosphorylation. However, we observed that IKK-deficiency had no detectable effect on starvation-induced ATG16L1 phosphorylation. (Fig. 2A, lanes 7-9). This is perhaps expected as IKKα has no known role in starvation-induced autophagy. This result indicates that the ATG16L1 subunit of the LC3-lipidating enzyme is a direct and physiological target of ULK under starvation.

We next asked if ULK contributed to ATG16L1 phosphorylation upon infection or if signalling went exclusively through IKKα. ULK wild-type, ULK dKO, or IKKα KO were infected with *Salmonella enterica* serovar Typhimurium (hereafter referred to as *Salmonella*) and ATG16L1 phosphorylation was examined by Western blot. Surprisingly, we observed that *Salmonella-*induced ATG16L1 phosphorylation was abolished in ULK dKO cells, but was still observed in IKKα knockout cells (Fig. 2B). These results clearly indicate that ULK is required for phosphorylation of ATG16L1 under both starvation and infection.

We next sought to determine the requirement for ULK and IKKα in promoting xenophagy. Xenophagic clearance of *Salmonella* is very well established and its intracellular growth is restricted by the pathway, making it an ideal model pathogen for this analysis. Wild-type or knockout cells were infected with *Salmonella* and the number of LC3B-positive *Salmonella* were quantified. LC3B is conjugated to the autophagosomal membrane and colocalizes with bacteria targeted for clearance by xenophagy and can be used at early time points to monitor xenophagy induction. We found that ULK-deficient cells exhibited a potent decrease in LC3B-positive bacteria, while IKKα loss did not significantly affect xenophagy (Fig. 2C, S2). In order to confirm the roles for ULK and IKKα in xenophagy induction and suppression of invasive bacteria we performed colony forming unit (CFU) assays in our wild-type or knockout lines. CFU assays measure bacterial viability after internalization and are inversely correlated with xenophagy rates^27^. Analysis of *Salmonella* viability 4 hours post infection revealed that ULK dKO cells harboured a much higher number of viable internalized bacteria, indicative of an autophagy defect, when compared to wild-type and IKKα knockout cells (Fig. 2D). Surprisingly, our results indicate that ULK, but not IKKα, is required for ATG16L1 phosphorylation and xenophagy induction.

Multiple groups have shown that the T300A substitution in caATG16L1 renders it sensitive to caspase cleavage under stress conditions including nutrient starvation and infection^21,24,28^. Moreover, it was shown that mutation of serine 278 of ATG16L1 to alanine protects against stress-induced caspase cleavage in the caATG16L1 background^24^. Our data indicate that ULK is responsible for the phosphorylation of wild-type ATG16L1 on S278 under nutrient starvation and infection. Therefore, we next sought to determine if ULK signalling was involved in the stress-induced destabilization of caATG16L1. HEK293A cells were transfected with either wild-type ATG16L1 or caATG16L1 co-transfected with increasing amounts of ULK kinase. Importantly, overexpression of ULK is known to result in autoactivation and induction of downstream signalling in the absence of stress, thereby allowing us to determine the isolated effect of ULK signalling on ATG16L1 stability independent of other stress-responsive pathways. Interestingly, we observed that ULK is capable of stimulating ATG16L1 cleavage and the level of cleavage is elevated in the caATG16L1 background (Fig. 3A). In order to determine if ATG16L1 cleavage was a result of ULK-mediated phosphorylation on S278 we transfected HEK293A cells with wild-type, T300A, or S278/T300A mutants of ATG16L1 in the presence or absence of ULK. Excitingly, we observed that single mutation of the ULK phosphorylation site was sufficient to reduce ULK-driven cleavage (Fig. 3B). These results indicate that caATG16L1 is preferentially cleaved through ULK-mediated phosphorylation of S278. Lastly, we repeated this experiment in the presence or absence of Z-VAD-FMK, a pan-caspase inhibitor, to confirm the faster migrating form of ATG16L1 was indeed a product of caspase-mediated cleavage. Treatment with a pan-caspase inhibitor resulted in a potent reduction in the levels of the faster migrating ATG16L1 band, confirming that the ULK-driven cleavage product was a caspase cleavage product (Fig. 3C). Increasing evidence *in vitro* and *in vivo* has shown that caspase-mediated destabilization of caATG16L1 is a critical event associated with the pathobiology of this SNP. Moreover, in unstressed conditions caATG16L1 is known to have the same stability as wildtype. However, the relationship between stress and the cleavage of caATG16L1 is not well understood. Collectively, our data shed light on the relationship between stress and caATG16L1 cleavage showing that: 1) ULK-mediated phosphorylation of ATG16L1 is increased under infection and starvation, which are known to promote the cleavage of caATG16L1, 2) caATG16L1 is preferentially cleaved upon ULK activation, and 3) mutating the ULK phosphorylation site reduces ULK-driven cleavage in the caATG16L1 background.

**Figure 3.**
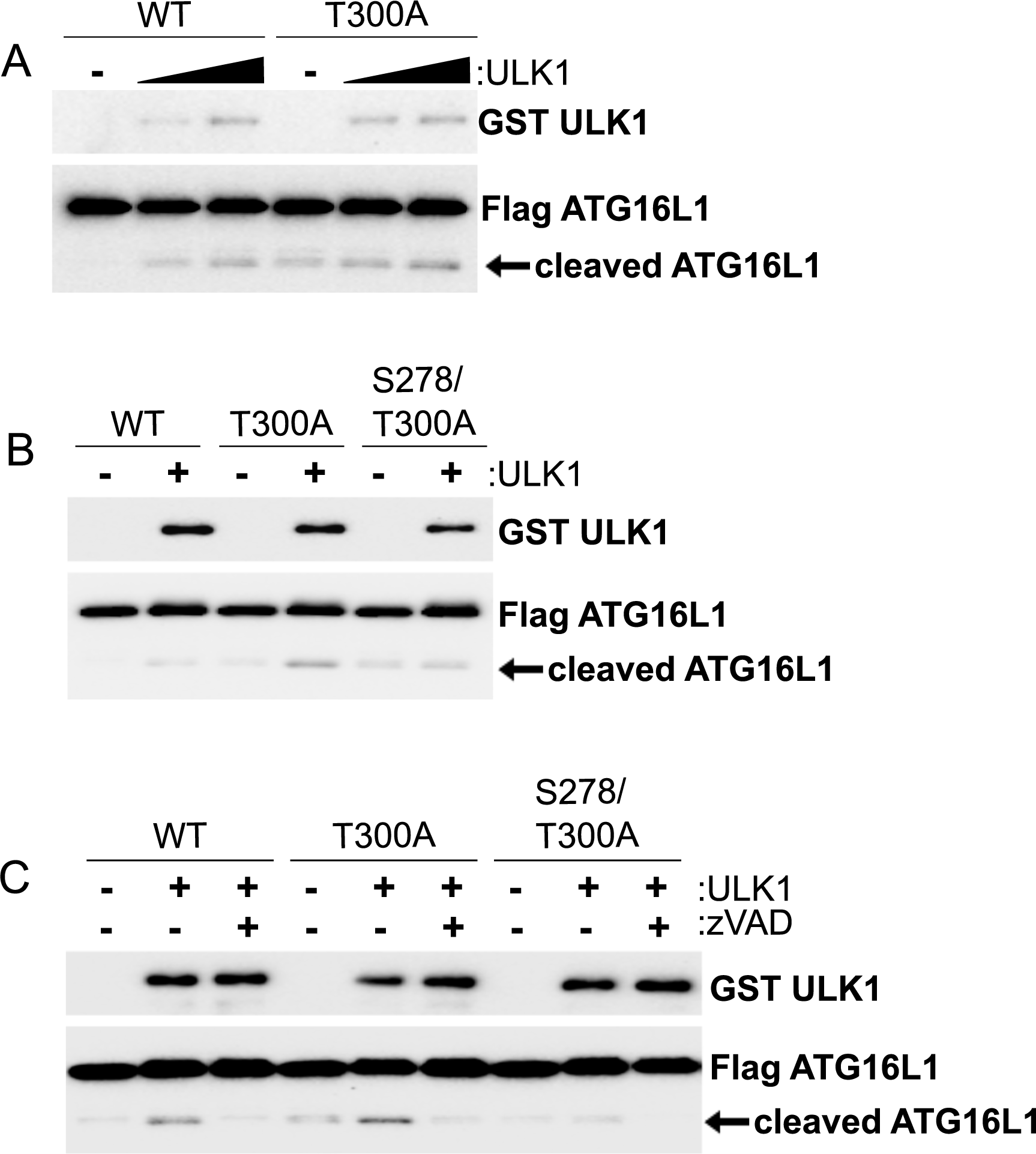
ULK promotes cleavage of caATG16L1 through phosphorylation on S278. **(A)** HEK293A cells were transfected with either tagged WT-ATG16L 1 or caATG16L1. ULK1 was co-transfected in increasing amounts where indicated. Cleavage of ATG16L1 was analyzed by WB of whole cell lysates. **(B)** HEK293A cells were transfected with either tagged wild-type, S278A, or S278/T300A ATG16L1 in the presence or absence of ULK. Cleavage of ATG16L1 was analyzed by WB. **(C)** HEK293A cells were transfected with the indicated plasmids in the presence or absence of Z-VAD-FMK (15 µM) for 4 hours. Cleavage of ATG16L1 was analyzed by WB.

ULK kinase has a well-established role in stimulating autophagy, making it unlikely that the primary function of ULK-induced ATG16L1 phosphorylation is to activate caspase-mediated cleavage. In order to identify the physiological role of ULK-mediated ATG16L1 phosphorylation we performed experiments on the wild-type protein, which is not cleaved as readily after phosphorylation. The best described function of ATG16L1 is to promote the correct localization of the E3-like enzyme that lipidates LC3 to the membrane of newly forming autophagosomes. Therefore, we first sought to determine if the localization of pATG16L1 differed from that of total ATG16L1 under infection. To compare localization we infected MEF with *Salmonella* and immunostained for bacterial lipopolysaccharide (LPS), pATG16L1, and total ATG16L1. We observed pATG16L1 primarily in the infected samples, confirming the reactivity of our antibody for IF (Fig. 4A). Excitingly, we found that pATG16L1 was preferentially localized with internalized bacteria (Fig. 4A). Analysis of total ATG16L1 staining also showed co-localization with bacteria, but the majority of the staining was diffuse in the cytoplasm (Fig. 4A). This could indicate that either ULK-mediated phosphorylation is important for ATG16L1 recruitment to bacteria, or that the phosphorylation occurs at the bacteria. We reasoned if phosphorylation of ATG16L1 affects bacterial localization then ULK-deficient cells should exhibit an impairment in ATG16L1 recruitment to pathogen. To test this hypothesis we infected wild-type or ULK-deficient cells and quantified the ability of total ATG16L1 to localize to internalized bacteria. Interestingly, we observed that the proportion of ATG16L1-positive bacteria in ULK-deficient MEF was reduced by over 80% compared to the wild-type controls (Fig. 4B, S3). Taken together our data indicate that ULK-mediated phosphorylation of ATG16L1 promotes localization to internalized bacteria.

**Figure 4.**
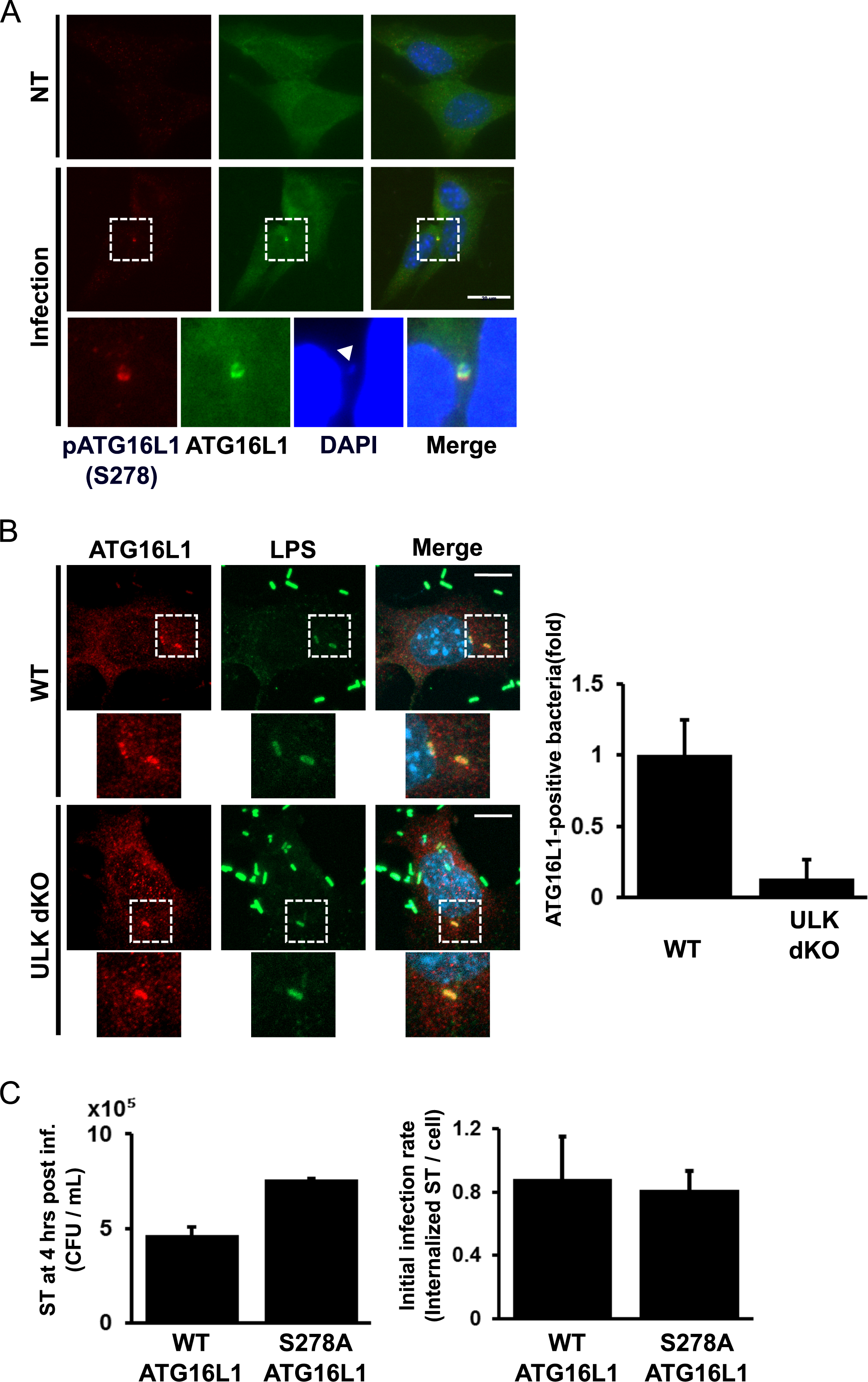
ULK-mediated phosphorylation is required for ATG16L 1 localization to Salmonella site and bacterial clearance. **(A)** Wild-type cells were infected with Salmonella for 1 hour. Phospho-ATG16L 1, total ATG16L 1, and DNA were stained and analysed by immunofluorescence. Representative immunofluorescent images are shown (scale bars, 20µm). White arrow highlights bacterial DNA. **(B)** Wild-type and ULK dKO were infected with Salmonella for 1 hour. lmmunofluorescence was performed using antibodies against LPS and ATG16L 1. Representative immunofluorescent images are shown on the left panel (scale bars, 10µm). Quantification of ATG16L 1-positive bacteria from 7 fields of view from a representative experiment is demonstrated on the right panel. **(C)** HCT116 stably expressing tagged wild-type or S278A ATG16L 1 were infected with ST for 1 hour. Xenophagy rates were examined through Colony Forming Unit (CFU) assays following removal of ST and incubation for 4 hours with gentamicin-containing media. Quantification of infection rates by immunofluorescence is shown in the right panel.

We next sought to determine the contribution of ATG16L1 phosphorylation to xenophagy. We stably reconstituted ATG16L1 KO HCT116 cells with either wildtype or the S278A mutant of ATG16L1 and performed a CFU assay. We found that the cells containing the phospho-dead mutant of ATG16L1 consistently had more *Salmonella* at 4 hours post-infection (Fig. 4C, left panel), despite being infected at the same rate (Fig. 4C, right panel). This indicates that ULK-mediated phosphorylation of wild-type ATG16L1 acts to promote xenophagy.

## Discussion

ULK has previously been described to phosphorylate several components of the autophagy-promoting lipid kinase complex to activate the autophagy pathway^10–12^. Here we have described that the autophagy E3-like enzyme is also regulated by ULK through direct phosphorylation of the ATG16L1 subunit. The discovery of a link between ULK and the LC3B-lipidating enzyme has raised several interesting lines of inquiry. For example, we have shown that wild-type ATG16L1 is also susceptible to ULK-sensitive caspase-mediated cleavage, albeit at a lower level than caATG16L1.

However, we currently do not know the physiological relationship between phosphorylation and caspase-mediated cleavage outside the context of the caATG16L1 allele. Potentially, caspase-mediated cleavage of ATG16L1 under stress represents a mechanism to curtail autophagy under severe or prolonged stress. Understanding the mechanistic link between apoptosis and autophagy may yield important conceptual advances.

Additionally, we have uncovered a role for ULK-signalling in CD through regulating the stability of caATG16L1. Interestingly, the functional significance of the S278 residue in CD had already been shown^24^. However, the lack of tools to measure endogenous pATG16L1 resulted in IKKα being identified as the kinase responsible for precipitating the phosphorylation and cleavage. Based on our data, as well as the previously reported link between starvation and pathogen-induced caATG16L1 dysfunction, we propose that ULK is the primary kinase responsible for ATG16L1 phosphorylation. However, it is quite possible that IKKα contributes to the destabilization of caATG16L1 through the previously reported activation of caspases^24^.

The preferential localization of pATG16L1 to internalized bacteria is also interesting. This is because frameshifts in the gene NOD2 are strongly associated with CD-development and have also been described to affect ATG16L1 localization to internalized bacteria^17^. This may imply a common defect of ATG16L1 function in CD. Consistent with this idea CD-associated SNPs have also been described in ULK1, albeit with less strength than ATG16L1 SNPs. As we have identified a functional redundancy between ULK1 and ULK2 in the promotion of ATG16L1 phosphorylation, which may explain the weak contribution of ULK1 polymorphisms in CD-susceptibility. Lastly, transcriptional repression of IRGM has also been linked to the development of CD. Molecularly, IRGM has been shown to bind both ULK1 and ATG16L1, although they have not been shown in a complex together. Therefore, it would be of value to determine if reductions in IRGM protein would have an effect on ULK-mediated ATG16L1 phosphorylation. Clearly, the identification of ULK-mediated ATG16L1 phosphorylation has opened up several avenues for future research, which will undoubtedly expand our understanding of xenophagy and the molecular basis of autophagy defects in CD.

## Supplemental Figure Legend

**Figure S1.**
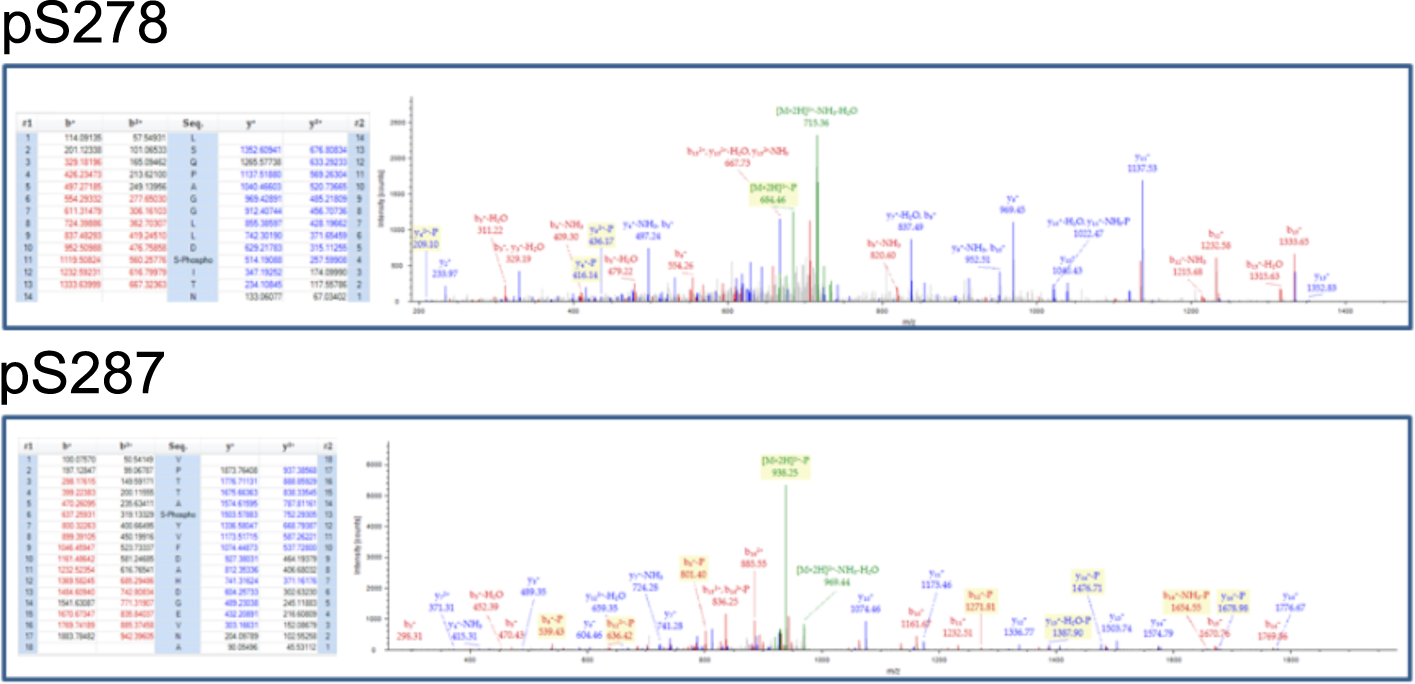
Mass spectrometry data of ATG16L 1 phosphorylation.

**Figure S2.**
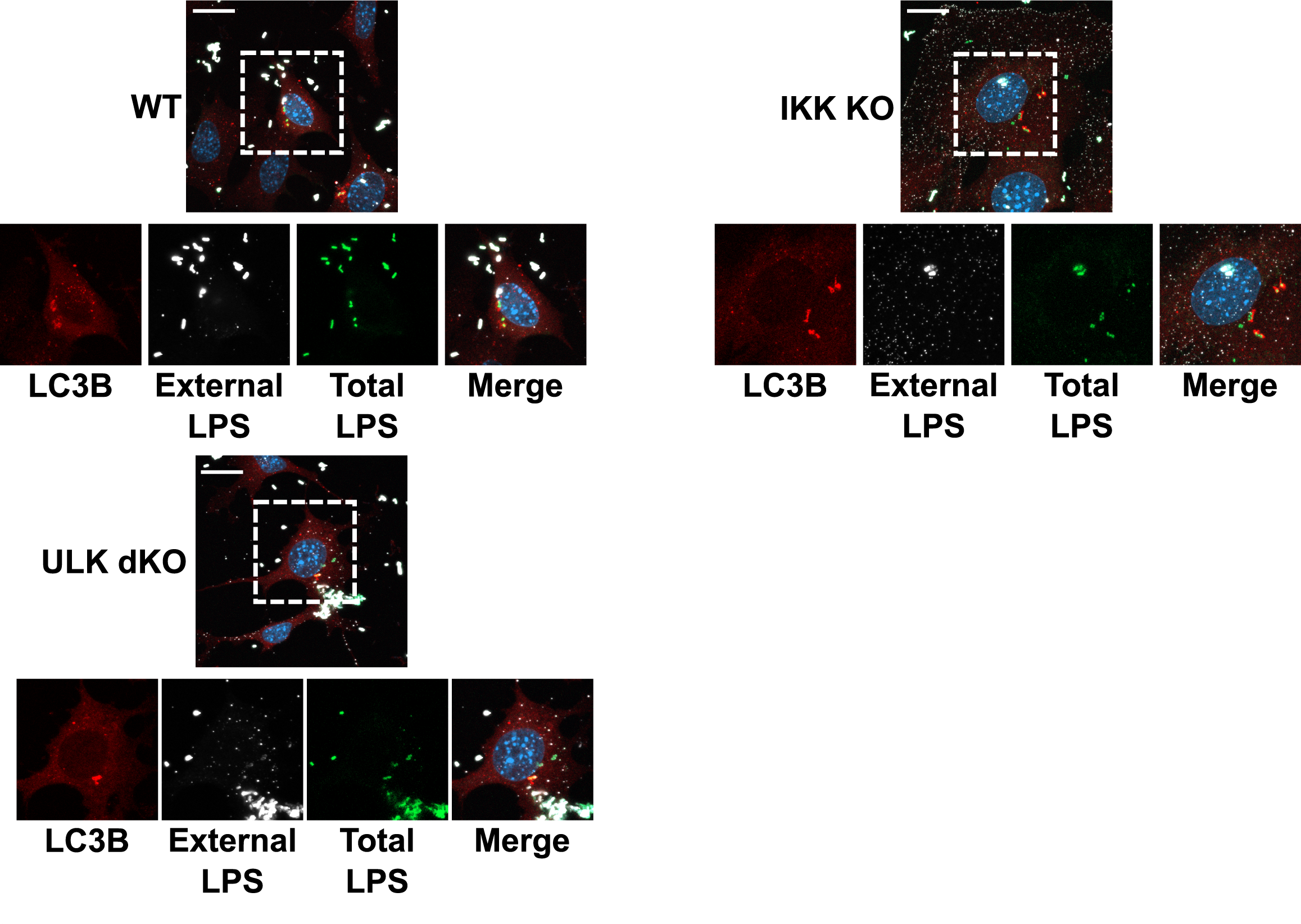
Enlargements of Fig. 2C with extracellular bacteria staining observable in white. MEF cells were were infected with Salmonella for 1 hour in the presence of Bafilomycin A 1. Endogenous LC38 (red) puncta was visualized (scale bars, 20µm) by immunofluorescence. Dashed boxes represent the cell selected for the main figure.

**Figure S3.**
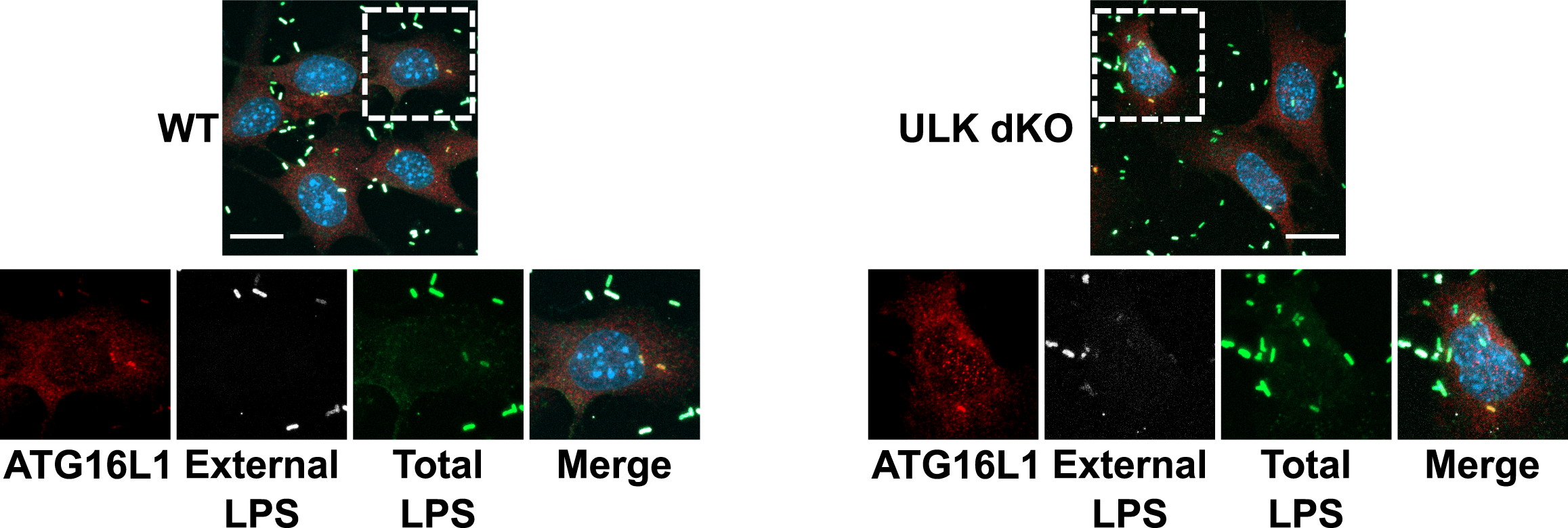
Enlargements of Fig. 4B with extracellular bacteria staining observable in white. MEF cells were were infected with Salmonella for 1 hour. Endogenous ATG16L 1 (red) puncta was visualized (scale bars, 20µm) by immunofluorescence. Dashed boxes represent the cell selected for the main figure.

## Material and Methods

### Antibodies and Reagents

Anti-IKKα (Cat#2682), HA-HRP (#cat 2999) and phospho-S6K T389 (Cat#9234) antibodies were obtained from Cell Signaling Technology. Anti-LC3B (Cat#PM036 for immunofluorescence) and ATG16L1 (Cat#PM040 for immunofluorescence) antibodies were purchased from MBL. Beta-Actin (Cat#A5441 clone AC-15) antibody was obtained from Sigma. DYKDDDDK Epitope Tag (Cat#NBP1-06712 for WB) antibody was purchased from Novus Biologicals. Anti-LPS FITC (Cat#sc-52223) and GST (Cat# sc-374171) antibodies was purchased from Santa Cruz Biotechnology. Anti-S6K (Cat#ab32529), LPS (Cat#ab128709), ATG16L1 (Cat#187671) antibodies were obtained from Abcam. phospho-ATG16L1 was made in collaboration with Abcam. Polyclonal sera was affinity purified by phospho peptide and recombinant ATG16L1 (non-phosphorylated) was mixed in at a 6:1 molar ratio (Rec. ATG16L1: IgG), prior to immunoblotting. Monoclonal phospho-antibody from a hybridoma generated from this rabbit was used for immunofluorescence (Abcam Cat#ab195242). Active GST-ULK1 (1-649) and GST-ULK2 (1-478) from insect cells were purchased from CQuential Solutions (Moraga, CA). Anti-His-HRP (Cat#460707) was obtained from Invitrogen. Z-VAD(OMe)-FMK (Cat#HY-16658-1MG) was purchased from MedChemExpress. Bafilomycin A1 was obtained from Tocris (Cat#133410U).

### Cell Culture

MEFs, HEK293A, and HCT116 were cultured in DMEM supplemented with 10% Bovine Calf Serum (VWR Life Science Seradigm). IKK wildtype and IKKα knockout cells were a generous gift from Dr. Michael Karin (University of California San Diego)^29^. ULK1/2 double knockout were a generous gift from Dr. Craig Thompson (Memorial Sloan Kettering)^30^. Amino acid starvation medium was prepared based on Gibco standard recipe omitting all amino acids and supplemented as above without addition of non-essential amino acids and substitution with dialyzed FBS (Invitrogen). Media was changed 1 hour before experiments.

### Generation of Stable Cell Lines

ATG16L1 knock-out lines were generated in the HCT116 background utilizing CRISPR/Cas9 targeting exon 1. The knock-out HCT116 clones were infected with retrovirus carrying either wild-type or phospho-dead ATG16L1 at different amounts in order to achieve near endogenous levels of ATG16L1.

### Bacterial Strains

Wild-type (SL1344) *Salmonella* was a gift from Dr. Subash Sad, (University of Ottawa). Bacteria were grown in Luria-Bertani broth (Fisher).

### Bacterial Infection

Salmonella were grown in 4 mL of LB broth at 37 degrees Celsius at 250 rpm. Overnight cultures of *Salmonella* were diluted 30-fold and grown until OD_600_ reached 1.5, followed by centrifugation of 10,000 g for 2 min, and resuspension in 1 mL of PBS. Bacterial stock was then diluted 5-fold (MOI = 900) in DMEM supplied with 10% heat-inactivated Bovine Calf Serum for infection. Cells cultured in antibiotic-free medium were infected with *Salmonella* and incubated at 37 degrees Celsius in 5% CO_2_ for the indicated time. Cells were washed in PBS once before direct lysis with 1X denaturing SDS sample buffer.

### Western Blot and Immunoprecipitation

Whole cell lysates were prepared by direct lysis with 1X SDS sample buffer. Samples were boiled for 10 min at 95 degrees Celsius and resolved by SDS-PAGE. Immune complexes were harvested from cells lysed in mild lysis buffer [10mM Tris pH 7.5, 10 mM EDTA, 100 mM NaCl, 50 mM NaF, 1% NP-40, supplemented simultaneously with protease and phosphatase inhibitor cocktails –EDTA (APExBIO)], followed by centrifugation at max speed for 10 minutes to remove cell debris. Protein A beads (Repligen) were washed 1X with PBS and incubated with antibodies and cell lysates for 1.5-3 hours followed by one 5-minute wash with MLB and inhibitors and 4 quick washes with MLB alone. Beads were boiled in 1X denaturing sample buffer for 10 min before resolving by SDS-PAGE.

### Site-Directed Mutagenesis

Primers used for T300A mutation are GGACAATGTGGATGCTCATCCTGGTTC (forward) and GAACCAGGATGAGCATCCACATTGTCC (reverse). Primers used for S278A mutation are GCCTTCTGGATGCTATCACTAATATC (forward) and GATATTAGTGATTGCATCCAGAAGGC (reverse). T300A followed by S278A mutation was performed to generate S278/T300A double mutation. Site-directed mutagenesis was performed based on KOD Xtreme Hot Start DNA Polymerase kit instructions purchased from Thermo Fisher. Specificity of mutagenesis was analysed by direct sequencing.

### Immunofluorescence

Cells were plated on IBDI-treated coverslips overnight. After treatments, cells were fixed by 4% paraformaldehyde in PBS for 15 min and subsequently permeabilized with 50 µg/mL digitonin in PBS for 10 min at room temperature. Cells were blocked in blocking buffer (1% BSA and 2% serum in PBS) for 30 min, followed by incubation with primary antibodies in the same buffer for one hour at room temperature. Samples were then washed 2X in PBS and 1X in blocking buffer before incubation with secondary antibodies one hour at room temperature. Slides were washed 3X in PBS, stained with DAPI, and mounted. Images were captured with inverted epifluorescent Zeiss AxioObserver.Z1. In the case of outside/inside bacterial staining, before permeabilization, the cells were incubated with anti-LPS antibody and corresponding secondary antibody in blocking buffer, accompanied by 3X PBS washes in between.

### Quantification of Immunofluorescence

An automated protocol built in the Image J software was used to analyse epifluorescent microscopy images to avoid bias. The same protocol was applied to each field of view and across samples. An average of 8 unique fields of view from representative experiments were selected for quantification.

### *in vitro* ULK Kinase Assay

HEK293A transiently expressing tagged ATG16L1 were immunoprecipitated. Pulldown proteins were washed 3X with MLB and 1X with MOPS buffer and were used as substrates for ULK kinase assay. ULK proteins were immunoprecipitated and extensively washed with MLB (once) and RIPA buffer (50 mM Tris at pH 7.5, 150 mM NaCl, 50 mM NaF, 1 mM EDTA, 1 mM EGTA, 1% SDS, 1% Triton X-100 and 0.5% deoxycholate) once, followed by washing with MLB buffer once followed by equilibration with ULK1 assay buffer (kinase base buffer supplemented with 0.05 mM DTT, 10 μM cold ATP, and 0.4 µCil ^32^P-ATP per reaction). Reactions were shaken at 250 rpm at 37 degrees Celsius for 30 min and stopped by direct addition of 4X sample buffer followed by 10 min boiling at 95 degrees Celsius and resolution by SDS-PAGE. The analysis of kinase reactions necessitated the separation of the kinase and substrate. *In vitro* kinase reactions were analyzed by autoradiograms.

### Colony Forming Unit (CFU) Assay

Cells were infected with *Salmonella* for 1 hour. The infected cells were washed 2X and incubated with media containing 100 µg/mL Gentamicin for 0.5 hour, followed by 4-hour incubation with media containing 50 µg/mL Gentamicin. The samples were rinsed 3X with PBS and lysed with CFU buffer (0.1% Triton X-100 and 0.01% SDS in PBS). The harvested lysates were serially diluted (1:100, 1:300, and 1:1000) and plated onto LB agar plates containing Streptomycin. The plates were incubated at 37 degrees Celsius for 16-18 hours and the colonies were counted to determine the number of CFU.

